# Fast and scalable off-target assessment for CRISPR guide RNAs using partial matches

**DOI:** 10.1101/2024.02.01.578509

**Authors:** Carl Schmitz, Jacob Bradford, Robert Salomone, Dimitri Perrin

## Abstract

The design of CRISPR-Cas9 guide RNAs is not trivial. In particular, it is crucial to evaluate the risk of unintended, off-target modifications, but this is computationally expensive. To avoid a brute-force approach where each guide RNA is compared against every possible CRISPR target site in the genome, we previously introduced Crackling, a guide RNA design tool that relies on exact matches over 4bp subsequences to approximate a neighbourhood and accelerate off-target scoring by greatly reducing the search space. While this was faster than other existing tools, it still generates large neighbourhoods. Here, we aim to further reduce the search space by requiring more, now non-contiguous, exact matches. The new implementation, called Crackling++, is benchmarked against our initial approach and other off-target evaluation tools. We show that it provides the fastest way to assess candidate guide RNAs. By using memory-mapped files, it also scales to the largest genomes. Crackling++ is available at https://github.com/bmds-lab/CracklingPlusPlus under the Berkeley Software Distribution (BSD) 3-Clause license.

## Introduction

CRISPR-based technologies have emerged as a very important tool for genome editing (1). At a high level, they rely on a CRISPR-associated (Cas) nuclease and an RNA sequence that ‘guides’ the nuclease to the targeted region. This guide RNA (gRNA) can be programmed to edit a specific region of the genome. In the case of Cas9, potential CRISPR target sites are defined by the presence a specific PAM (NGG), and the 20 bases upstream from it are used to construct the gRNA. Over the past decade, scientists have used CRISPR to enable both fundamental research and wide-ranging applications such as developing animal models of disease (2), studying endangered species (3), increasing crop resilience (4), and even developing new therapies (5).

However, while the range of applications demonstrate the flexibility of this technology, designing gRNAs is not a trivial task. In particular, it is crucial to limit the risk of edits at genomic locations other than the targeted one. Here, using a 20bp sequence that is unique to this region is of course necessary, but it is not sufficient. A CRISPR site that has a different but similar-enough 20bp sequence can still be incorrectly targeted. This is known as an off-target modification. An off-target site is typically defined as having 1 to 4 singlebase mismatches compared to the intended one.

Various methods have been proposed to evaluate that off-target risk, such as the Zhang score (6) and the Cutting Frequency Determination (CFD) score (7). Both methods use a similar strategy: they calculate a score for the candidate gRNA against any off-target site, referred to as a *local score*, and aggregate this over all possible off-target sites for that gRNA, thus generating a *global score* that represents the specificity of that gRNA. They differ in how they calculate that local score. The Zhang score is based on the position of the mismatches between the gRNA and the off-target site, while the CFD score considers the position and type of mismatch. Details, including all equations needed to calculate these scores, are given in (8). However, what they share, and what is also common to other methods, is that they require a list of all off-target sites for that gRNA.

In other words, for each gRNA, it is necessary to identify across the *entire* genome *every* CRISPR site that is at most 4 mismatches away from that gRNA. A brute-force approach calculating all pairwise distances would scale quadratically with the number of CRISPR sites in the genome. This makes it unpractical, as large genomes contain billions of such sites. To avoid this brute-force approach, we introduced a gRNA design tool called Crackling (8). It employs a data structured called Inverted Signature Slice Lists (ISSL), originally developed for searching web-scale collections of data (9), which we use as an approximate look-up table for identifying candidate off-target sites. These lists rapidly extract a *neighbourhood* for any candidate gRNA, such that: (i) all the actual off-target sites for that gRNA are contained in the neighbourhood, and (ii) the neighbourhood is much smaller than the entire list of CRISPR sites for the whole genome. The first property ensures that we can calculate exact Zhang or CFD scores (as no genuine off-target site is missed). The second property provides the speed-up compared to the brute-force approach.

ISSL arranges the search space so the neighbourhood can be extracted in a constant time, and is constructed as follows. First, each site is split into contiguous slices of a fixed equal length. In (8), we used five non-overlapping slices, each of them four nucleotides long. For example, if the site is AATTCTGATTGGACGTTGCA, then the associated slices are: slice one – AATT, slice two – CTGA, slice three – TTGG, slice four – ACGT, and slice five – TGCA. The number corresponding to each slice is called its *position*.

Following this, a hash map is constructed using a combined key of the position and nucleotide sequence. Each key associates a list of CRISPR sites referred to as a *partial neighbourhood*. All members of that partial neighbourhood have the same nucleotide sequence in the specified position (see Fig.2 in (8) for a diagrammatic example). The overall neighbourhood for a given guide is obtained by retrieving the partial neighbourhood for each slice.

Importantly, using this five-slice configuration ensures that all genuine off-target sites will share *at least* one slice with the candidate guide. If a CRISPR site differs from the candidate guide on all five slices, then by construction there will be at least five mismatches between them. In that case, this means that the site is not an off-target site for that guide, and we do not need it in the neighbourhood for the guide. Constructing the neighbourhood using slice identity therefore achieves both properties listed above.

ISSL uses bit-encoded representations of the sites to reduce the amount of memory required and more efficiently utilise CPU instructions. A two-bit encoding is used to represent each nucleotide in the genomic alphabet (A as 00, C as 01, G as 10, T as 11). Given CRISPR sites are 20-nucleotide sequences, 40 bits (of a uint64 object) are needed for the encoding.

When calculating the off-target scores for a candidate guide, the guide is prepared using the same encoding and slicing strategy as the sites. For each slice, the members of the associated neighbourhood are compared to the candidate guide using the Hamming distance to identify off-target sites. This is necessary because the ISSL approach does not make any guarantees on the tightness of the neighbourhood. If the distance is four or less, the site is confirmed as a genuine offtarget site, and the local off-target score is calculated. Importantly, each site is considered only once per candidate guide, despite the opportunity of it being identified as a candidate off-target site via multiple slices.

We showed that Crackling provided an order of magnitude speedup compared to other tools such as Cas-OFFinder (10), Crisflash (11), and FlashFry (12).

Here, our focus is on redesigning the neighbourhood construction so that we can generate tighter ones while still guaranteeing that no genuine off-target site is missed. We will compare our new results with Crackling and with these other tools. We will also include a newer method called CRISPRSE, which was reported to be faster than FlashFry and CrisFlash (13).

## Materials and Methods

### A. Reducing the neighbourhood size

Crackling achieved a significant speedup because it does not need to consider all possible CRISPR sites. As described above, it extracts for each candidate guide a neighbourhood that contains all the genuine off-target sites for this guide, but is smaller than the entire set of sites.

However, while membership of that neighbourhood is a *necessary* condition for a site to be a genuine off-target site, it is not a *sufficient* one. They are many sites that are more than 4 mismatches away from the guide (and are therefore not actual off-target sites for this guide) but still are an exact match to the guide on a given 4bp slice. This leads to very large neighbourhoods, and unnecessary computations.

The maximum size of the partial neighbourhood retrieved from each slice depends on the length of that slice. Crackling uses 5 slices of length 4. This means that for each slice, there are 16 positions (20 − 4) that are left ‘free’ in any CRISPR site retrieved by that slice. As each of these positions can assume any of A, T, C and G, the theoretical limit for that partial neighbourhood is 4^16^ (4,294,967,296) elements. In practice, for any genome and any candidate guide, not all of these alternatives will exist, but the neighbourhoods can still be large.

Requiring longer matches automatically reduces the size of the neighbourhood. For instance, for a slice of length 5, the maximum size comes down to 4^15^ (1,073,741,824). For a slice of length 8, the maximum size is only 65,536 (4^8^).

However, increasing the slice length breaks the first property we aim to achieve for the neighbourhood: even at a length of 5, we are already no longer guaranteed that all off-target sites are included in the neighbourhood. For instance, if we consider AATTCTGATTGGACGTTGCA as the candidate guide, it is possible to have off-target sites that do not share any 5bp subsequence, such as AATTGTGATAGGAGGTTGGA (where the mismatches are shown in red). However, the two sequences still share multiple sets of 5 positions on which they match. To reduce the neighbourhood size, we need to move from contiguous matches, i.e. *slices*, to non-contiguous ones, which we will refer to as *masks*.

We also need to consider strategies to construct sets of masks that will capture all possible off-target sites.

### B. Valid sets of masks

By construction, a set of masks satisfies the second property we want for our neighbourhood: it is necessarily smaller than the entire list of CRISPR sites for the whole genome. We consider that the set is valid if it also satisfies the first property: all the actual off-target sites for that gRNA are contained in the neighbourhood.

This will be achieved when, for any possible off-target site, there is at least one mask in the set over which the gRNA and that off-target site are an exact match. We say that the mask *captures* this off-target site.

We can reflect about this in an abstract way in terms of the position of the mismatches between a guide and an offtarget site, independently from the sequences themselves. We will use a 20-bit binary representation where 1 indicates a match and 0 indicates a mismatch. For example, the off-target combination 01111111111111111110 indicates that the off-target site and the candidate guide have mismatches at the first and last positions. If the gRNA is AATTCTGATTGGACGTTGCA, then this notation represents off-target sites such as CATTCTGATTGGACGTTGCT, GATTCTGATTGGACGTTGCC, etc.

Similarly, we can use a binary representation for the mask, where 1 indicates a position where the mask requires a match and 0 indicates a position that the mask ignores. We say that a mask has a weight *x* if it is has exactly *x* bits equal to 1. For instance, 11100000000000000011 is the mask of weight 5 that considers the first three and last two positions.

We can use a logical AND operator between an off-target combination and a mask. The mask captures the offtarget combination *if and only if* the operator returns a sequence where exactly *x* bits are equal to 1. For instance, the mask above would capture the off-target combination 11110111011011111011, but it would not capture 11110111011011111110.

It is worth noting that, if *x* ≤ 16 then for any off-target combination of 4 mismatches (or less), there is always at least one mask of weight *x* that captures that combination.

### C. Constructing an initial valid set

In practice, our focus will be on ensuring that the set captures all combinations of exactly 4 mismatches anywhere in the sequence. There are 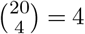 such combinations. If it does, then it also necessarily captures all the combinations of 1-3 mismatches. Let *A* be the list of all combinations of 4 mismatches.

Next, we created a list *B* of all possible masks of weight *x*. There are 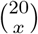 such masks. For instance, for *x* = 8, there are 125, 970 possible masks.

Our objective is to construct a valid set *C* of masks. As defined above, it is a set that captures all possible off-target combinations.

We start from an empty set *C*. We iterate through *B*, and for each mask *m* ∈ *B*, we count how many elements from *A* it can capture. The mask *m*^*^ that captures the highest number of elements is added to *C*. It is also removed from *B*, and *all* the combinations that it captured are removed from *A*. We repeat that process until *A* is empty. Because we know that each configuration can always be captured by at least one mask, |*A*| is decreasing by at least 1 in each iteration. This guarantees that the algorithm terminates in a finite number of iterations and that, when it does, *C* captures all combinations. Thus, *C* is a valid set.

However, while this greedy approach ensures that *C* is a valid set, it makes no guarantee about its size. In fact, |*C*| can be quite large. This can become a problem for large genomes. For any given genome, the index built using *C* maintains one reference per mask for each site. Increasing the number of masks increases the number of references per site, and therefore the memory footprint of the program (which will be larger than that of the initial slice-based implementation of Crackling).

To offset the increased memory requirement, the index can be mapped to virtual memory space, avoiding the limit of physical memory. However, reading from memory-mapped files is slower due to the decreased input-output bandwidth that secondary storage has compared to main memory, so it is still desirable to minimise |*C*|. Index size and the impact of memory mapping are discussed in detail in Section J.

### D. Set optimisation

Imagine that we want to check whether it is possible to generate a valid set where |*C*| = *y*. We know that there are 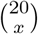 masks of weight *x*. For *x* = 8, we get 125, 970 masks of weight *x*. we get 125,970 possible masks, and the greedy approach produces a set such that |*C*| = 28. If we aim for *y* = 27, the number of possible sets is given by:

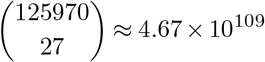

For *x* = 10, there are 184, 756 possible masks, and the greedy approach provides |*C*| = 46. If we aim for *y* = 45, the number of possible sets is given by:

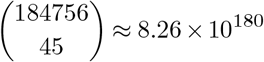

Clearly, a brute-force approach would not work at these scales, and we have developed an approximate search instead. We employed a variant of the adaptive splitting method for optimisation (14) to complete the combinatorial optimisation to reduce the number of masks.

To search for a valid set containing *y* masks, we started by randomly generating a population of sets of weight *x*. Each set was then individually scored based on the number of offtarget combinations it captures, and the population of sets was ranked accordingly. The goal is to find a set that captures all off-target configurations, i.e. a valid set, giving it a perfect score.

If no such set has been identified yet, the top 20% of the sets were selected and randomly used to rebuild a new population. Each set in the new population was passed through a Markov Monte Carlo mutation algorithm. The mutation function guarantees that the new set will detect more offtarget configurations (or at least as many) than the mask configuration passed into the function. By repeating this process of mutating the top-scoring sets in each population, the sets get increasingly better until the algorithm is stopped or a valid set is found.

Once a valid set has been found for a value of *y*, the optimisation process is repeated for *y* − 1, until no valid set can be found in a reasonable time.

### E. Benchmark setup

We benchmarked the performance of our new mask-based method against the initial slice-based Crackling implementation, all the tools reported in (8), and CRISPR-SE.

We used all genomes from the previous study, as well as four larger ones that existing tools could not process. All were obtained from the National Center for Biotechnology Information (NCBI) Genome database, using the accession numbers given in Table 1.

**Table 1.** Benchmark genomes obtained from the NCBI genome database.

| Genome | Accession Number | Size (Mb*) | Off-target Count | Unique Off-target Count |
| --- | --- | --- | --- | --- |
| *Oryza sativa* | GCF_001433935.1 | 374.4 | $61.7 \times 10^6$ | $44.3 \times 10^6$ |
| *Panicum hallii* | GCF_002211085.1 | 507.4 | $97 \times 10^6$ | $51.2 \times 10^6$ |
| *Xiphophorus couchianus* | GCF_001444195.1 | 685.5 | $101.1 \times 10^6$ | $89.1 \times 10^6$ |
| *Parus major* | GCF_001522545.3 | 998.3 | $177.1 \times 10^6$ | $171.8 \times 10^6$ |
| *Crotalus viridis* | GCA_003400415.2 | 1297 | $201.2 \times 10^6$ | $158.8 \times 10^6$ |
| *Danio rerio* | GCF_000002035.6 | 1679.4 | $214.8 \times 10^6$ | $121 \times 10^6$ |
| *Oncorhynchus mykiss* | GCF_002163495.1 | 1950 | $499.9 \times 10^6$ | $181.6 \times 10^6$ |
| *Mus musculus* | GCF_000001635.24 | 2730.7 | $588.6 \times 10^6$ | $368.9 \times 10^6$ |
| *Monodelphis domestica* | GCF_000002295.2 | 3502.4 | $326.1 \times 10^6$ | $441.7 \times 10^6$ |
| *Triticum urartu* | GCA_003073215.1 | 4661.6 | $862.4 \times 10^6$ | $212.2 \times 10^6$ |
| *Rhinatrema bivittatum* | GCF_901001135.1 | 5178.6 | $1012.1 \times 10^6$ | $583 \times 10^6$ |
| *Triticum turgidum* | GCA_900231445.1 | 9964.3 | $1763.2 \times 10^6$ | $448 \times 10^6$ |
| *Agapanthus africanus* | GCA_030015575.1 | 20168.2 | $2929.3 \times 10^6$ | $369.1 \times 10^6$ |
| *Ambystoma mexicanum* | GCA_002915635.3 | 28206.9 | $5379.9 \times 10^6$ | $2660.2 \times 10^6$ |
| *Neoceratodus forsteri* | GCA_016271365.1 | 34557.6 | $1363.1 \times 10^6$ | $794 \times 10^6$ |
| *Protopterus annectens* | GCF_019279795.1 | 40054.3 | $6299.3 \times 10^6$ | $2639.7 \times 10^6$ |

\* Megabases

As in (8), the sites used for the off-target scoring includes both canonical CRISPR sites as well as non-canonical ones ending with NAG rather than the standard NGG.

All the files required by CRISPR-SE were created as per the published instructions (13). All other tools are used as previously described (8). For our new mask-based implementation, we tested four configurations with different values for *x* and |*C*|.

Each tool was benchmarked by averaging the time taken to process 5 unique sets of 10,000 randomly selected guides on genomes of increasing size. The computer used for the benchmark was the same as the one used by (8), a Linux workstation with one 8-core Intel Core i7-5960X (3.0 GHz) CPU, 32 GB RAM, 32 GB allocated swap space, and a Samsung PM87 SSD.

## Results

### F. Set size

We tested our mask-based solution for two mask weights, *x* = 8 and *x* = 10. As guaranteed by its structure, the greedy approach was able to generate valid sets for both. For *x* = 8, that ‘greedy set’ contained |*C*| = 28 masks. For *x* = 10, |*C*| = 46.

We therefore started the optimisation process described in Section D with *y* = 27 for *x* = 8, and *y* = 45 for *x* = 10. The optimisation was able to identify valid sets with |*C*| = 20 and |*C*| = 37 for the two mask weights, respectively. It is possible that smaller valid sets exist. Our focus here is on establishing the overall principles and testing the resulting configurations, rather than on guaranteeing the optimality of the set.

It is important to note that the optimisation process is completely independent from any genome the method could process: the sets are valid for any genome, and are computed only once. It was a one-off task that no user of the tool would have to handle (unless they want to try to optimise further).

In what follows, we tested five mask configurations:

- 5 masks of weight 4, which are constructed to match the 5 slices of length 4 from the original Crackling implementation. This is noted 4–5.
- 28 masks of weight 8, obtained from the greedy approach. This is noted 8–28.
- 20 masks of weight 8, from the optimisation; noted 8– 20.
- 46 masks of weight 10, from the greedy approach; noted 10–46.
- 37 masks of weight 10, from the optimisation; noted 10–37.

### G. Total run time

Table 2 presents the total off-target scoring run time for Crackling++ (in the 8–20 mask configuration), CRISPR-SE and the results published in (8). The offtarget score reported by Crackling++ and other tools using the Zhang and/or MIT scores is between 0 and 100 (or between 0 and 1), where a higher score means a lower risk of off-target modifications. In Crackling++, we can set a threshold *τ* on this off-target to decide whether to accept or reject a candidate gRNA. As this global off-target score depends on the sum of the local scores, it is possible to terminate the calculation as soon as the score is compared to be below *τ*, as discussed in (8). Here, we report results for *τ* = 75 (a conservative value which limits the risk of off-target modifications whilst still accepts several guides per gene, see e.g. (15)) and *τ* = 0 (which effectively guarantees that the execution continues until the final off-target score is calculated).

**Table 2.** Algorithm run times.

| Genome | fM | fMB | FlashFry | Crisflash | Cas-OFFinder | Crackling | CRISPR-SE* | Crackling++<br>($\tau = 0$) | Crackling++<br>($\tau = 75$) |
| --- | --- | --- | --- | --- | --- | --- | --- | --- | --- |
| Oryza sativa | 46:17 | 10:46 | 02:24 | 13:04 | 01:26 | 00:28 | 00:29 | 00:13 | 00:11 |
| Panicum hallii | 140:49 | 11:12 | 02:30 | 13:43 | 01:24 | 00:43 | 00:33 | 00:15 | 00:20 |
| Xiphophorus couchianus | 315:20 | 22:24 | 04:56 | 16:09 | 01:33 | 01:17 | 00:43 | 00:24 | 00:20 |
| Parus major | 134:29 | 33:05 | 07:19 | — | 02:06 | 02:25 | 00:54 | 00:40 | 00:24 |
| Crotalus viridis | 309:51 | 26:42 | 05:52 | — | 01:53 | 02:13 | 00:53 | 00:38 | 00:19 |
| Danio rerio | 178:37 | 23:37 | 05:12 | — | 01:45 | 01:31 | 00:47 | 00:31 | 00:14 |
| Oncorhynchus mykiss | 240:19 | 30:16 | 06:50 | — | 02:00 | 02:25 | 00:59 | 00:41 | 00:16 |
| Mus musculus | 383:51 | 41:52 | 09:18 | — | — | 05:13 | 01:06 | 01:20 | 00:19 |
| Monodelphis domestica | 500:43 | 36:04 | 07:58 | — | — | 06:10 | 01:06 | 01:38 | 00:20 |
| Triticum urartu | 612:32 | 306:15 | 05:03 | — | — | 03:48 | 01:08 | 00:45 | 00:09 |
| Rhinatrema bivittatum | — | — | — | — | — | — | — | 01:53 | 00:26 |
| Triticum turgidum | — | — | — | — | — | — | — | 01:34 | 00:19 |
| Agapanthus africanus | — | — | — | — | — | — | — | 01:25 | 00:16 |
| Ambystoma mexicanum | — | — | — | — | — | — | — | 05:17 | 01:57 |
| Neoceratodus forsteri | — | — | — | — | — | — | — | 02:22 | 00:35 |
| Protopterus annectens | — | — | — | — | — | — | — | 06:11 | 01:53 |

Time in (MM:SS) format
fM is findMatches. fMB is findMatchesBit (forked version of fM which has been adapted to use bit-encoded sequences).
— indicates runtime failure due to memory limitations
\* indicates that than an algorithm only searched for off-target sites and did not produce any scores.

When considering only the ten genomes used in (8), from *O. sativa* (0.37 gigabases) to *T. urartu* (4.7 gigabases), Crackling++ with *τ* = 75 is the fastest tool across all genomes. When running with *τ* = 0, Crackling++ was still the fastest tool on eight of these genomes, with CRISPR-SE faster on the other two. However, CRISPR-SE only searches for offtarget sites and does not provide any off-target scores, which would then have to be calculated separately.

The CRISPR-SE article provides instructions for parsing its output to the formats needed by CRISPOR (16) and FlashFry (12), so that these tools can use CRISPR-SE’s output to calculate off-target scores. Here, we did not attempt this, as Crackling++ with *τ* = 75 was demonstrated to be faster and more scalable than CRISPR-SE alone, even before attempting to calculate those scores.

Crackling++ was the only tool to process all genomes in the benchmark, up to and including the largest ones, while the other tools were ended by the operating system’s out-of-memory killer. To our understanding, there is no other CRISPR off-target scoring tool that can process the largest genomes in the NCBI Genome database.

These results are also shown in Figure 1. Here, the horizontal axis uses the number of unique off-target sites in the genome. This better represents the search space, and therefore the problem size for the measured algorithms, rather than directly using the genome size.

**Fig. 1.**
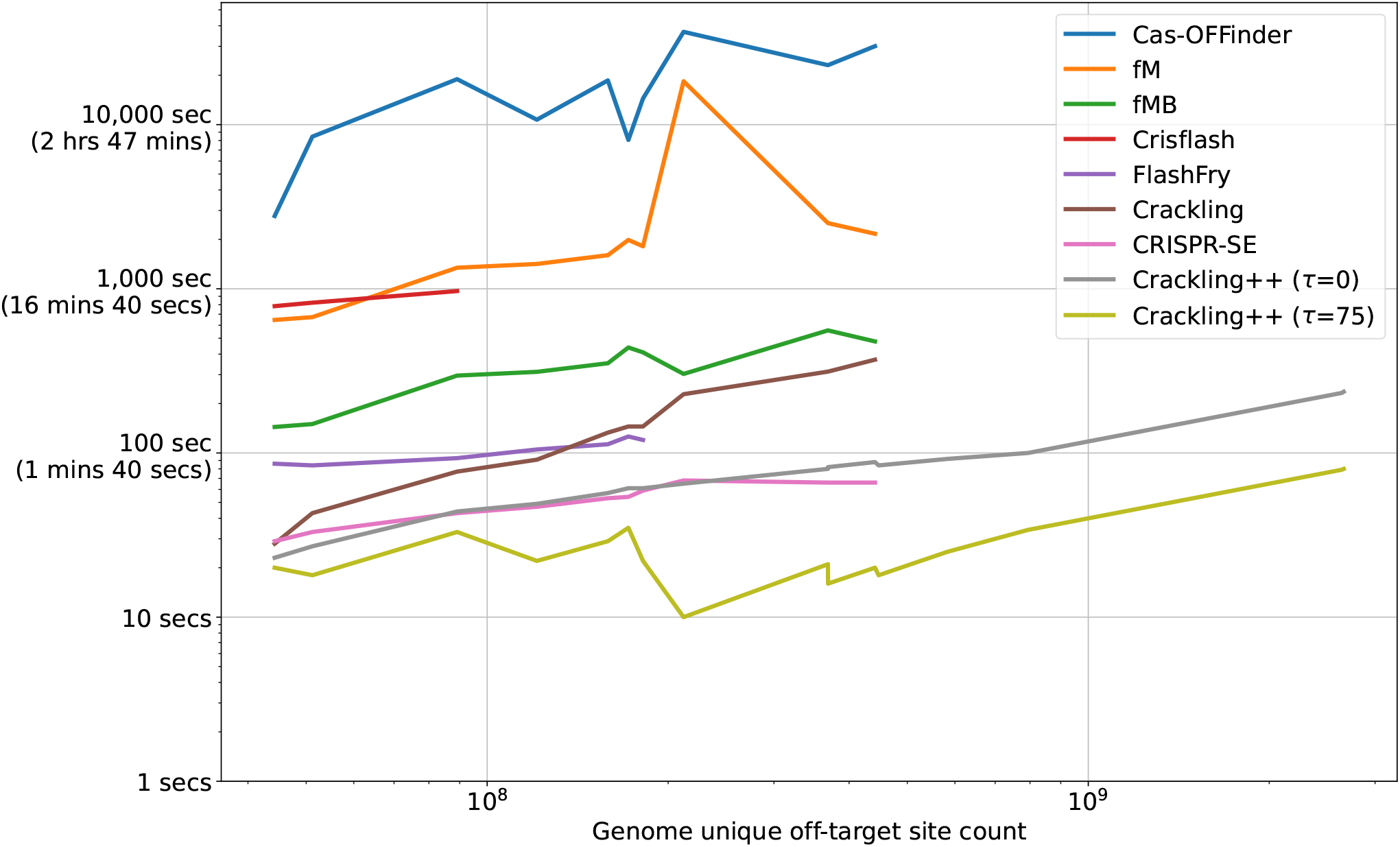
Run times of different tools based on the number of unique off-target sites in the genome

### H. Run time of alternative mask configurations

Tables 3 and 4 show the loading and processing times of different Crackling++ slice configurations for score *τ* = 75 and *τ* = 0, respectively. The first important result is that, for all genomes and for all configurations of masks and thresholds, the memory-mapped implementation is faster than the in-memory implementation. This is the result of the balance between loading time and processing time. For each indisvidual guide, the processing is faster using the in-memory implementation. This is because retrieving a neighbourhood in memory is faster than doing so on disk. However, that implementation first needs to load the whole index in memory. This takes more time as the genome size increases, as shown in Table 5. It quickly outgrows the small gains from the inmemory scoring. Overall, this makes the memory-mapped implementation faster, and the difference between the two implementations increases with the genome size.

**Table 3.** Total run times for Crackling++ with different configurations. Results are for a threshold equal to 75.

| Genome | 4-5 IM | 8-28 IM | 8-20 IM | 10-46 IM | 10-37 IM | 4-5 MM | 8-28 MM | 8-20 MM | 10-46 MM | 10-37 MM |
| --- | --- | --- | --- | --- | --- | --- | --- | --- | --- | --- |
| *Oryza sativa* | 00:15 | 00:23 | 00:17 | 00:38 | 00:32 | 00:14 | 00:16 | 00:11 | 00:24 | 00:20 |
| *Panicum hallii* | 00:14 | 00:27 | 00:20 | 00:44 | 00:36 | 00:12 | 00:14 | 00:11 | 00:23 | 00:18 |
| *Xiphophorus couchianus* | 00:36 | 00:50 | 00:36 | 05:28 | 01:04 | 00:33 | 00:27 | 00:20 | 00:42 | 00:33 |
| *Parus major* | 00:50 | 06:48 | 01:10 | — | 03:53 | 00:43 | 00:29 | 00:24 | 00:37 | 00:35 |
| *Crotalus viridis* | 00:41 | 06:17 | 01:04 | — | 07:39 | 00:34 | 00:24 | 00:19 | 00:31 | 00:29 |
| *Danio rerio* | 00:29 | 01:05 | 00:47 | 07:43 | 05:43 | 00:23 | 00:18 | 00:14 | 00:25 | 00:22 |
| *Oncorhynchus mykiss* | 00:39 | 07:15 | 01:13 | — | — | 00:31 | 00:19 | 00:16 | 00:24 | 00:22 |
| *Mus musculus* | 01:05 | — | — | — | — | 00:45 | 00:22 | 00:19 | 00:20 | 00:21 |
| *Monodelphis domestica* | 01:12 | — | — | — | — | 00:48 | 00:21 | 00:20 | 00:20 | 00:20 |
| *Triticum urartu* | 00:31 | 08:27 | 05:41 | — | — | 00:19 | 00:09 | 00:09 | 00:10 | 00:10 |
| *Rhinatrema bivittatum* | 01:27 | — | — | — | — | 00:47 | 00:26 | 00:26 | 00:25 | 00:25 |
| *Triticum turgidum* | 01:03 | — | — | — | — | 00:28 | 00:19 | 00:19 | 00:18 | 00:18 |
| *Agapanthus africanus* | 00:55 | — | — | — | — | 00:31 | 00:16 | 00:16 | 00:16 | 00:16 |
| *Ambystoma mexicanum* | — | — | — | — | — | 04:11 | 02:06 | 01:57 | 01:45 | 01:20 |
| *Neoceratodus forsteri* | 07:31 | — | — | — | — | 01:04 | 00:36 | 00:35 | 00:34 | 00:34 |
| *Protopterus annectens* | — | — | — | — | — | 03:51 | 01:58 | 01:53 | 01:38 | 01:19 |

IM is in-memory. MM is memory mapped.
— Indicates runtime failure due to memory limitations

**Table 4.** Total run times for Crackling++ with different configurations. Results are for a threshold equal to 0.

| Genome | 4-5 IM | 8-28 IM | 8-20 IM | 10-46 IM | 10-37 IM | 4-5 MM | 8-28 MM | 8-20 MM | 10-46 MM | 10-37 MM |
| --- | --- | --- | --- | --- | --- | --- | --- | --- | --- | --- |
| *Oryza sativa* | 00:26 | 00:25 | 00:17 | 00:39 | 00:32 | 00:24 | 00:18 | 00:13 | 00:29 | 00:23 |
| *Panicum hallii* | 00:30 | 00:28 | 00:21 | 00:44 | 00:37 | 00:27 | 00:20 | 00:15 | 00:33 | 00:27 |
| *Xiphophorus couchianus* | 00:56 | 00:50 | 00:37 | 05:45 | 01:04 | 00:51 | 00:34 | 00:24 | 00:52 | 00:44 |
| *Parus major* | 01:48 | 07:12 | 01:11 | — | 09:14 | 01:40 | 00:55 | 00:40 | 01:16 | 01:01 |
| *Crotalus viridis* | 01:40 | 06:39 | 01:06 | — | 07:46 | 01:32 | 00:52 | 00:38 | 01:13 | 00:57 |
| *Danio rerio* | 01:15 | 01:08 | 00:49 | 07:56 | 05:44 | 01:10 | 00:43 | 00:31 | 01:01 | 00:49 |
| *Oncorhynchus mykiss* | 01:48 | 07:57 | 01:16 | — | — | 01:39 | 00:57 | 00:41 | 01:17 | 01:01 |
| *Mus musculus* | 03:42 | — | — | — | — | 03:24 | 02:00 | 01:20 | 01:39 | 01:20 |
| *Monodelphis domestica* | 04:37 | — | — | — | — | 04:13 | 02:25 | 01:38 | 01:50 | 01:28 |
| *Triticum urartu* | 01:56 | 09:19 | 06:02 | — | — | 01:46 | 01:05 | 00:45 | 01:24 | 01:05 |
| *Rhinatrema bivittatum* | 05:32 | — | — | — | — | 05:03 | 02:46 | 01:53 | 01:54 | 01:32 |
| *Triticum turgidum* | 03:58 | — | — | — | — | 03:38 | 02:17 | 01:34 | 01:46 | 01:24 |
| *Agapanthus africanus* | 03:23 | — | — | — | — | 03:07 | 02:01 | 01:25 | 01:45 | 01:22 |
| *Ambystoma mexicanum* | — | — | — | — | — | 453:29 | 07:04 | 05:17 | 05:06 | 03:56 |
| *Neoceratodus forsteri* | 43:03 | — | — | — | — | 10:48 | 03:23 | 02:22 | 02:04 | 01:40 |
| *Protopterus annectens* | — | — | — | — | — | 476:07 | 08:23 | 06:11 | 05:29 | 03:52 |

IM is in-memory. MM is memory mapped.
— Indicates runtime failure due to memory limitations

**Table 5.**
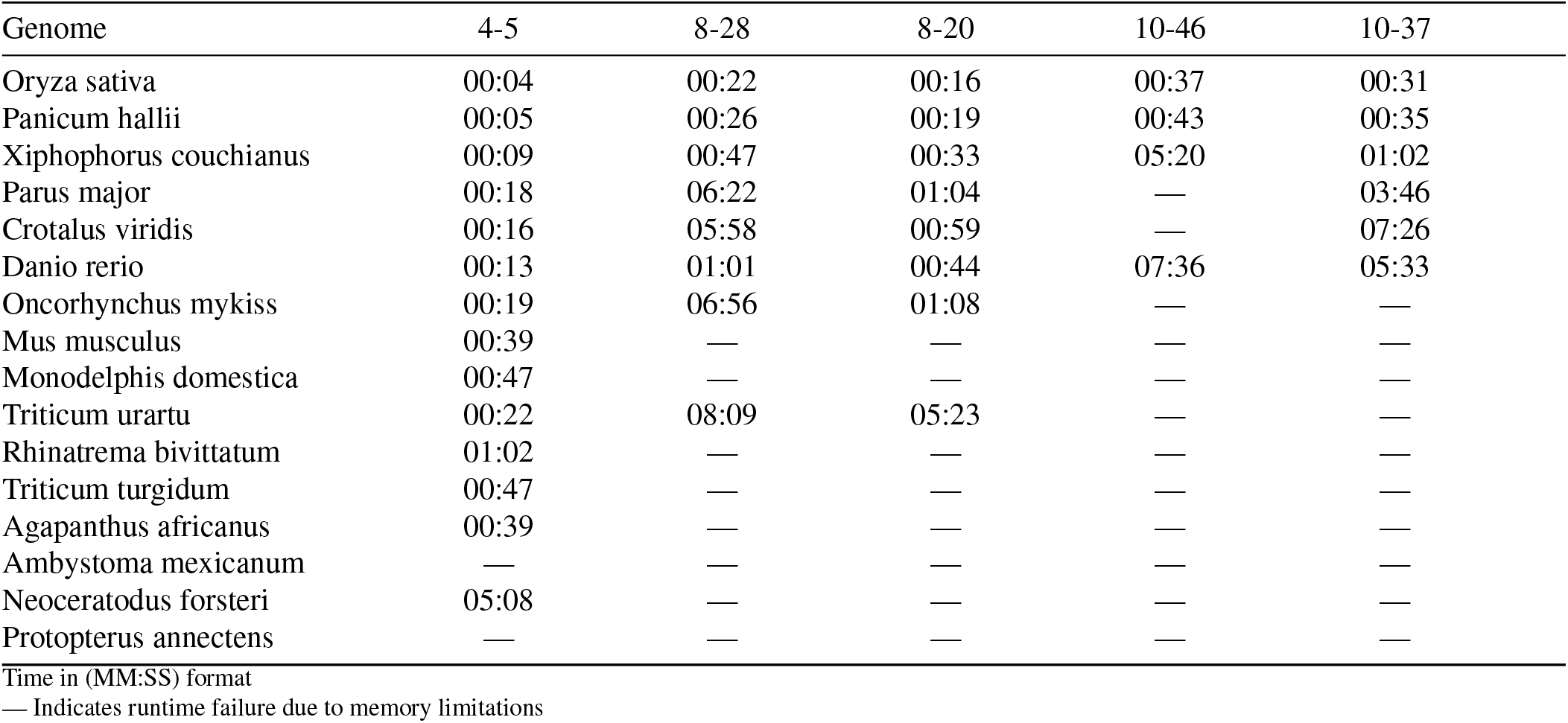
Loading times for different in-memory Crackling++ configurations.

The memory-mapped implementation is also the only one able to successfully process all genomes, irrespective of their size. This is a direct result of removing the need to load the index in memory.

For a given mask weight, the implementation using the optimised mask set is always faster than the implementation using the set generated with the greed approach. This is entirely expected: once the mask weight is fixed, neighbourhoods will on average have similar sizes, but with a smaller set we process fewer. The difference is more visible for *τ* = 0 (which means processing all sites until the score is final), but it is still present for *τ* = 75.

For large genomes (e.g. axolotl, Australian lungfish), the best results are obtained for a mask weight *x* = 10, which retrieves the smallest neighbourhoods. For small genomes (e.g. rice, zebrafish), neighbourhoods tend to always be small (as there are not as many CRISPR sites overall), and the best results are for *x* = 8, which has a smaller *number* of neighbourhoods. For medium-sized genomes (e.g. mouse), there is no clear difference between the two configurations.

## Discussion

### I. Neighbourhood reduction drives faster off-target assessment

As we had hypothesised, the speed-up in Crackling++ is obtained thanks to a great reduction in the number of sites that are processed. This is shown in Figure 2. By increasing the weight of the masks (from 4 to 8, and then to 10), we reduce the number of ‘free’ positions in the sequence, and therefore reduce the size of the neighbourhood that each mask will extract. That Figure also explains why, for each mask weight, the optimised set gives better results than the set obtained with the greedy approach: fewer masks means fewer neighbourhoods, and therefore fewer sites to process.

**Fig. 2.**
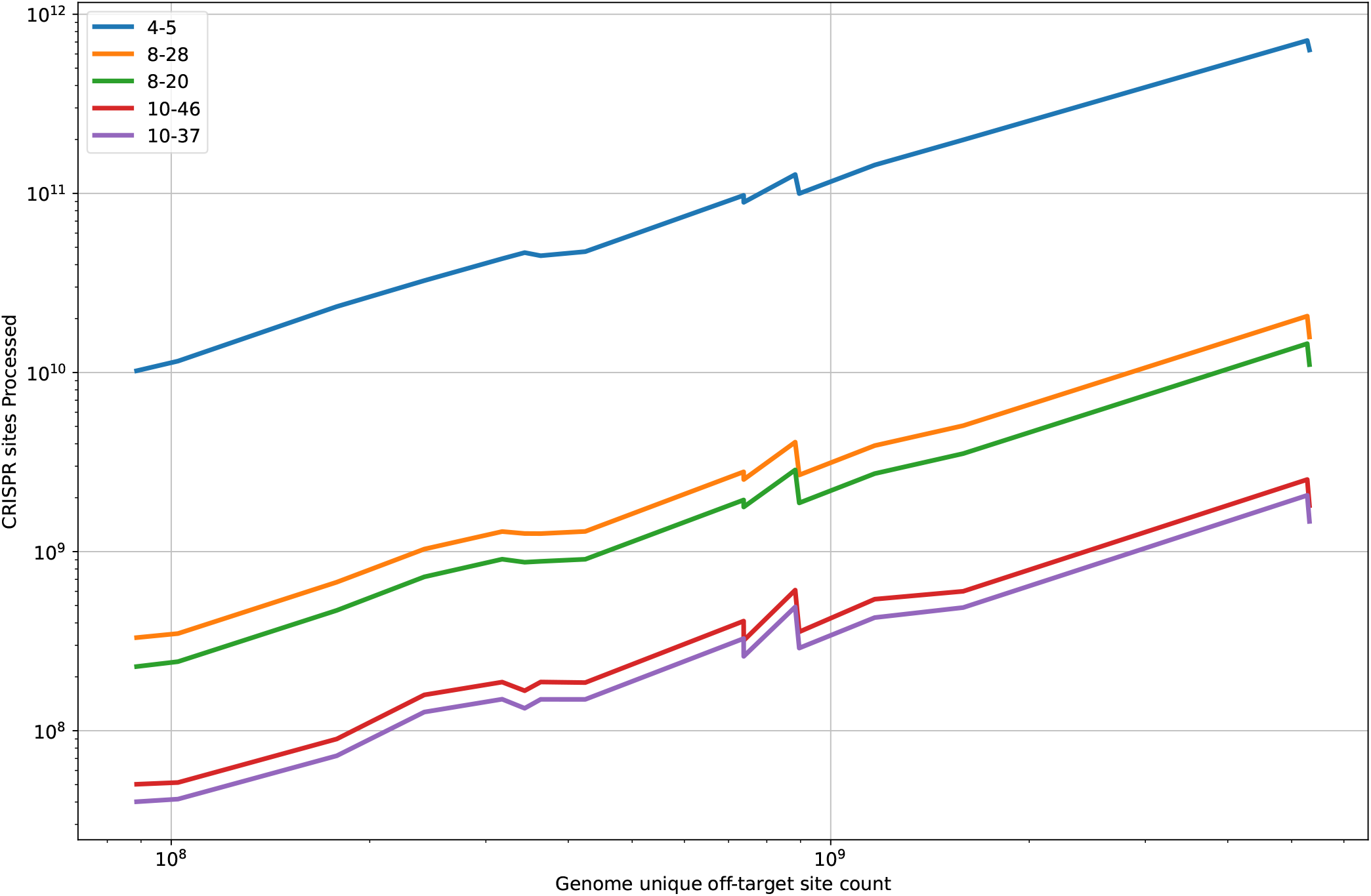
Total number of CRISPR sites processed by each mask configuration

It is important to also remember that, for small genomes, fewer comparisons (with *x* = 10) did not mean faster execution (compared to *x* = 8). As we discussed in the previous Section, the effective size of the neighbourhoods was already small enough. However, for these genomes, the run time is short enough under all configurations that the value chosen for *x* does not matter.

Overall, our recommendation is to use the default configuration (10–37 with *τ* = 75) for all genomes.

### J. Increased memory footprint can be addressed with memory mapping

As previously described, the size of the index is directly impacted by |*C*|, the size of the mask set: each mask independently splits the list of CRISPR sites into neighbourhoods, and more masks therefore translates into a larger index. When we increase the mask weights *x*, more masks are needed to cover all possible off-target combinations, and |*C*| increases.

We measured the empirical size of the indexes for each of the mask configurations presented in this paper. Compared to the configuration in (8), the index was on average 4.84× larger for the 8–28 configuration, 3.5× larger for 8–20, 7.89× larger for 10–46, and 6.37× larger for 10–37. Similarly, the index data structure used in CRISPR-SE was 4.67× larger than the index for the original 4–5 configuration of Crackling.

This increases the size of the overall index to the point that, for larger genomes, a standard workstation generally would not have enough available memory to load the index in its entirety. This is evidenced by the fact that CRISPR-SE could not process genomes larger than *T. urartu* (see Table 2), and that the in-memory implementation of Crackling++ stops working earlier when |*C*| increases (see ‘IM’ columns in Tables 3 and 4). We also see that, even before reaching the size where the out-of-memory killer is triggered, large genomes can saturate the physical RAM and access swap memory, which leads to degraded performance (Table 5).

Not only does an increased index size limit the portability of the tool on small-memory machines, but it also increases loading time for all machines, as shown in Table 5. To mitigate these two issues, we introduced the memory-mapped implementation: we no longer load the entire index into main memory, but instead map it to virtual memory. The ‘MM’ columns in Tables 3 and 4 show that this implementation can process all genome sizes, and that it is faster than the in-memory version (as it removes the loading times reported in Table 5).

Reading from virtual memory space is slower as data is retrieved directly from disk, bounding speed to the input– output bandwidth of the disk, but the loading time for the inmemory implementation increases faster than this difference. Unless scoring a very large number of guides on a genome small enough for the index to fit in memory, the memory-mapped implementation will always be preferable.

Crackling++ provides both implementations, but we recommend using the memory-mapped version as per the default configuration.

### K. Future directions

The memory-mapped implementation for the mask-based extraction of neighbourhoods provides a fast and scalable approach to off-target scoring for genomes of any size. Crackling++ was able to process even the largest known genomes.

However, while that speed-up is obtained through a reduction in the neighbourhood size, these neighbourhoods are not perfect. They still contain a large number of sequences that are more than four mismatches away from the guide RNA being scored. This is shown in Figure 3. The proportion of true off-target sites is increasing with larger mask weights. For a given mask weight, it is also increasing with smaller mask sets. Yet, it still remains below 10% across all genomes and all configurations. It means that the vast majority of sequences are still too far from the guide to contribute to the off-target score: they are discarded and generate unnecessary computation.

**Fig. 3.**
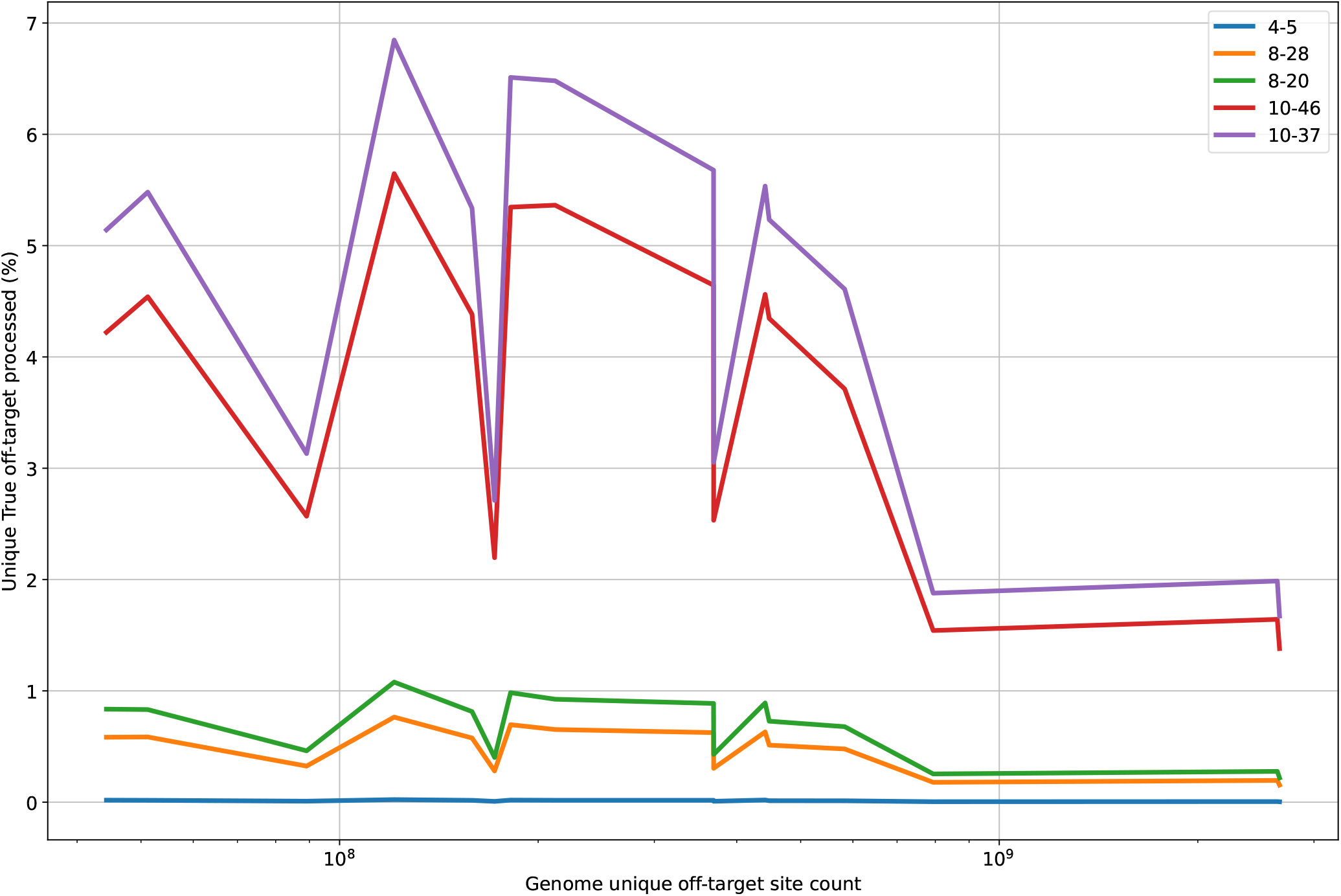
Comparing the percentage of unique true off-targets processed by each mask configuration

In theory, this means that higher mask weights could provide further improvements. However, in practice, this is probably not going to be feasible for large genomes. Larger values for *x* would necessarily mean larger values for |*C*| as well. Because each mask in *C* distributes all sites across neighbourhoods, the resulting index size would be impractical. The memory-mapped implementation is addressing the memory footprint, but not the disk storage requirement that extreme values for |*C*| would induce. Future improvements to off-target scoring are likely to rely on fundamentally different approaches.

Crackling++ is built with these considerations in mind. It is a complete re-implementation in C++ of the original Crackling pipeline (which was largely Python-based apart from the off-target scoring component). It follows a modular design, so that each module can be maintained, updated or replaced independently, reducing the amount of work required to update the tool.

The index used for scoring can be built for any valid mask set *C*. The only requirement is a file containing all the binary masks for that set. This allows the users to explore different mask configurations, including through further optimisation or different mask weights. For convenience, the five configurations tested in this article (4–5, 8–28, 8–20, 10–46 and 10–37) are available in the tool repository.

## Conclusions

In this paper, we investigated the use of partial matches to accelerate the off-target scoring step in the design of CRISPR guide RNAs. We show that, by moving from small contiguous slices to long non-contiguous masks as the basis for neighbourhood extraction, we can reduce the number of sites that need to be evaluated during off-target scoring. This directly leads to a significant speed-up of that scoring.

We also introduced an optimisation process to find small valid sets for any given mask weight. A smaller set means a faster execution, and the process also guarantees the validity of the set (and therefore that the off-scoring remains exact). To offset the increased memory required by the mask-based method (compared to short slices), we also provided and validated an approach that utilises memory-mapped files to use indexes that do not fit in physical memory. This allows the tool to scale to genomes of any size on consumer hardware. We benchmarked this new tool, Crackling++, on a range of configurations and genome sizes, and showed that it is a fast and scalable solution that outperforms other existing tools. Crackling++ is available at https://github.com/bmdslab/CracklingPlusPlus under the Berkeley Software Distribution (BSD) 3-Clause license.

## ACKNOWLEDGEMENTS

C.S. is supported by an Australian Government Research Training Program Scholarship. D.P. is supported by the Australian Research Council (ARC Discovery Project DP210103401).

## Notes

### Competing Interest Statement

The authors have declared no competing interest.

